# White matter microstructure is associated with the precision of visual working memory

**DOI:** 10.1101/2023.01.23.525278

**Authors:** Xuqian Li, Dragan Rangelov, Jason B. Mattingley, Lena Oestreich, Delphine Lévy-Bencheton, Michael J. O’Sullivan

## Abstract

Visual working memory is critical for goal-directed behaviour as it maintains continuity between previous and current visual input. Functional neuroimaging studies have shown that visual working memory relies on communication between distributed brain regions, which implies an important role for long-range white matter connections in visual working memory performance. Here, we characterised the relationship between the microstructure of white matter association tracts and the precision of visual working memory representations. To that purpose, we devised a delayed estimation task which required participants to reproduce visual features along a continuous scale. A sample of 80 healthy adults performed the task and underwent diffusion-weighted MRI. We applied mixture distribution modelling to quantify the precision of working memory representations and guess rates, both of which contribute to observed responses. Latent components of tract-specific microstructural indices were identified by principal component analysis. Higher working memory precision was associated with lower bulk diffusion across ten tracts of interest and higher directionality of diffusion in a group of frontoparietal-occipital tracts. Importantly, there was no association between guess rates and any of the structural components. Our findings suggest that microstructural properties of white matter tracts connecting posterior and frontal brain regions mediate, in a functionally specific manner, the precision of visual working memory.

## 1. Introduction

Visual working memory involves active maintenance and manipulation of visual information over a short period of time. It supports a range of cognitive functions and contributes to goal-directed behaviour (Awh & Jonides, 2001; de Fockert et al., 2001; Gathercole et al., 2004; Henderson et al., 2014). Functional magnetic resonance imaging (fMRI) and positron emission tomography (PET) studies have shown that visual working memory relies on a widespread network of brain regions including the lateral prefrontal cortex, anterior insula, posterior parietal cortex, inferior temporal cortex, and early visual cortex (Daniel et al., 2016; Owen et al., 2005; Rottschy et al., 2012; Wager & Smith, 2003). Recent studies using network-based approaches have shown increased global efficiency and decreased modularity of functional networks during working memory compared with rest, potentially supporting coordinated neural activity (Dagenbach, 2019). The idea that visual working memory relies on communication across a large-scale network implies a critical role for long-range white matter connections that support information transmission in the brain (Düzel et al., 2010; Pajevic et al., 2014). Here, we characterised the relationship between white matter microstructure and visual working memory performance in a large sample of neurotypical adult humans.

Previous studies have related visual working memory performance to white matter microstructure in several long-range association tracts (Lazar, 2017). In particular, the superior longitudinal fasciculus (SLF), a major pathway that connects the parietal and frontal lobes (Makris et al., 2005), has been related to visual working memory performance (Darki & Klingberg, 2015; Vestergaard et al., 2011). The SLF can be further divided into dorsal (SLF I), middle (SLF II), and ventral components (SLF III), with different patterns of connectivity to other brain regions and, arguably, distinct functional specialization (Makris et al., 2005; Parlatini et al., 2017; Thiebaut de Schotten et al., 2011). Further, the inferior frontal-occipital fasciculus (IFOF), a tract that mediates direct communication between occipital and frontal lobes (Forkel et al., 2014), has also been associated with visual working memory (Krogsrud et al., 2018; Peters et al., 2014; Walsh et al., 2011). Finally, the inferior longitudinal fasciculus (ILF), a temporal-occipital tract that runs more superficially and ventrally than the IFOF (Herbet et al., 2018), also plays a role in modulating visual working memory function, based on evidence from healthy children and case studies of focal lesions (Krogsrud et al., 2018; Shinoura et al., 2007). In the current study, we focused on the relationship between visual working memory performance and five critical tracts-of-interest (TOIs): the SLF I, SLF II, SLF III, IFOF and ILF.

Conceptually, participant responses in visual working memory tasks can reflect a true but noisy memory representations of target items or random guesses originating from attentional lapses, poor task compliance or other factors. Unfortunately, previous investigations that aimed to relate visual working memory performance to white matter microstructure did not distinguish these sources of behavioural variability (Darki & Klingberg, 2015; Krogsrud et al., 2018; Nagy et al., 2004; Peters et al., 2014; Vestergaard et al., 2011; Walsh et al., 2011). The overarching goal of the current study was to use computational modelling of behaviour to independently characterise contributions of white matter microstructure to these theoretically distinct components of visual working memory performance.

To evaluate visual working memory performance, we developed a novel version of the delayed estimation paradigm (Emrich et al., 2013; Gorgoraptis et al., 2011; Taylor & Bays, 2020), which required participants to encode three visual gratings that varied in both their spatial location and orientation. After a short delay period, participants reproduced, on a continuous scale, either the location or orientation of only one of the gratings, as indicated by a probe which appeared after the memory maintenance period. The different features to be retrieved (location and orientation) were included to test the extent to which brain-behaviour associations are similar across the spatial and non-spatial domains of visual working memory. Having participants respond on a continuous scale allowed us to calculate error magnitudes that could be modelled using mixture distribution modelling to independently estimate *the precision* of “true” working memory responses and *the proportion of responses arising from random guessing* (Bays et al., 2009; Zhang & Luck, 2008). To decrease dimensionality of the white matter microstructural data, we used a two-step principal component analysis (PCA) to estimate latent components over different *measures* and different *tracts*. If the microstructure of long-range white matter tracts selectively affects the precision of visual working memory representations, then the structural components should correlate exclusively with response precision but not with random guess rates. Additionally, if the brain-behaviour relationships are common for different features in visual working memory, there should be no difference between the location and orientation tasks for any of the observed associations.

## 2. Methods

### 2.1. Participants

Eighty-seven healthy adult volunteers were recruited from The University of Queensland through an online volunteer system. Seven participants were excluded from subsequent analyses due to data corruption (*n* = 4) or incomplete MRI data (*n* = 3). The final sample included 80 participants aged from 18 to 38 years (*M* = 24.24, *SD* = 4.61; 39 females). All participants completed safety screening questionnaires and provided written informed consent before the experimental sessions. Participants were reimbursed at a rate of $20 per hour. The study was approved by the Human Research Ethics Committee of The University of Queensland.

### 2.2. Experiment

#### 2.2.1. Apparatus

Stimuli were displayed on an LCD monitor (VG248QE) with a resolution of 1920×1080 pixels and a refresh rate of 60 Hz. Participants were seated approximately 60 cm from the monitor in a dimly illuminated room, with their head position maintained with a chinrest. The experiment was implemented under MATLAB R2018a (MathWorks, Natick, MA) using Psychtoolbox (Brainard, 1997; Pelli, 1997). Eye position was recorded using a desk-mounted eye-tracking system sampled at 120 Hz (iView RED-m infrared eye tracker, SensoMotoric Instruments, Teltow, Germany). The eye-tracker was calibrated and validated before each experimental block using a five-point calibration grid. The experiment was performed concurrently with electroencephalography (EEG) recording but the EEG data are not reported in this manuscript.

#### 2.2.2. Visual Working Memory Experiment

The location and orientation versions of the visual working memory tasks were presented in separate blocks in random order for each participant (Fig. 1). In each trial, participants were first presented with a black central cross (size: 1.32 degrees in visual angle [dva]; RGB: 0, 0, 0; line thickness: 0.13 dva) and a black arrow (RGB: 0, 0, 0; width: 2.64 dva; height: 0.66 dva) located 2.64 dva above the fixation for 300 ms, reminding the participants to encode only items on the left or right side of the screen. The “Cue” display was followed by the “Encoding” display presented for 400 ms and comprising six oriented gratings (radius: 2.64 dva). The gratings were randomly arranged on an invisible circle (radius: 10.55 dva) with respect to their centre. Any two adjacent gratings were separated by at least 20° and maximally 90° centre-to-centre offset and no grating was presented within ±15° range from the vertical midline. The grating orientations were randomly sampled over a 0°-179° range in increments of 2°. Following the encoding display, the “Maintenance” display appeared for 900 ms, comprising only the central fixation. Next, depending on the task, a “Probe” display showing either the location or orientation of one of the memorized gratings on the cued side was presented for 700 ms. Participants were instructed to maintain fixation until the “Response” display. If an eye movement or blink was detected, a trial was discarded and replayed at the end of the block.

**Figure 1.**
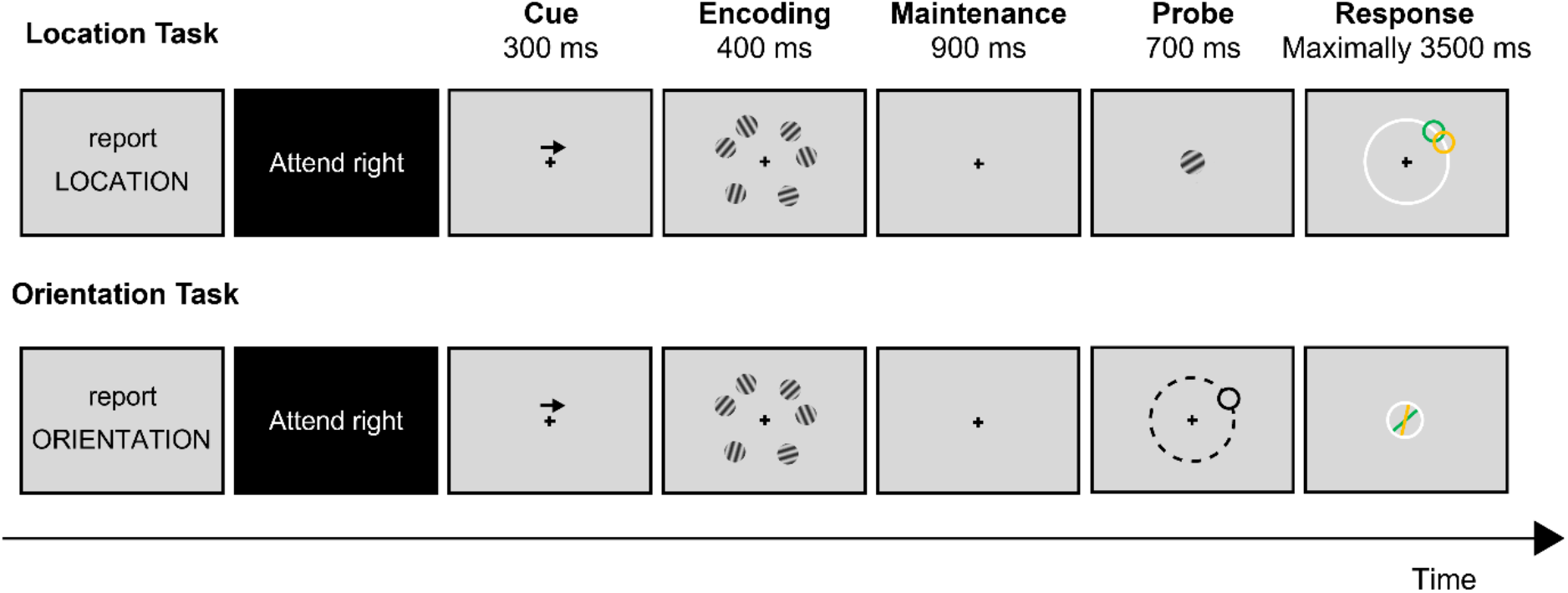
Schematic Illustration of the Location and Orientation Tasks. At the start of each block, an instruction message appeared to indicate the subsequent task. At the beginning of each run, a message appeared on a black background to remind participants to encode items presented on either the left side or right side. In each trial, six gratings were presented during the encoding period. After a 900 ms delay period, participants provided a response based on the probe information.

For the location task, one of the memorized gratings on the cued side was presented in the centre of the probe display. This probe item displayed the orientation of one of the three presented gratings on the cued side, independently of its location. Simultaneously with probe disappearance, a white response circle appeared in the “Response” display in the middle of the screen (radius: 10.55 dva; RGB: 255, 255, 255; line thickness: 0.13 dva). Participants reported the location of the target grating on the response circle using a computer mouse. At the beginning of the response display, the mouse cursor was set to the centre of the screen. Immediately after participants moved the cursor, a smaller black circle (radius: 2.64 dva; RGB: 0, 0, 0) was revealed with a red dot (radius: 0.66 dva; RGB: 255, 0, 0) placed at its centre. While moving the mouse, both circle and red dot were locked to the cursor movement to allow participants to adjust their response. A maximum of 3500 ms was set for the response period. Once a response was made, or at the end of the response period, a green feedback circle (radius: 2.64 dva; RGB: 0, 255, 0) representing the correct location of the probed item was shown for 1000 ms.

For the orientation task, a black circle outline (radius: 2.64 dva; RGB: 0, 0, 0) corresponding to the exact location of one of the three gratings on the cued side was presented in the probe display. This probe item displayed the spatial location of one of the three presented gratings on the cued side, independently of its orientation. Simultaneously with probe disappearance, a white response circle appeared in the “Response” display in the middle of the screen (radius: 2.64 dva; RGB: 255, 255, 255; line thickness: 0.13 dva). Participants reported the orientation of the target grating using the mouse. Similar to the location task, the mouse cursor was set to the centre of the screen. As soon as participants started moving the cursor, a yellow line (RGB: 255, 255, 0; line thickness: 0.13 dva) inside the response circle was revealed and locked to the cursor movement. This helped the participants to adjust their response within a maximum response period of 3500 ms. Immediately following the response, or at the end of the response period, a green feedback line (RGB: 0, 255, 0; line thickness: 0.13 dva) representing the correct orientation was shown for 1000 ms.

Prior to the main experiment, all participants completed one practice block for each task with four trials per cued side (2 tasks × 2 cues × 4 trials = 16 trials). The main experiment consisted of four randomized blocks so that each of the tasks was presented twice. Each block contained two runs in which participants were cued to encode items on the left side of the screen, and two runs in which participants were cued to encode items on the right side, and these alternated with each other. The runs were counterbalanced across participants, with some participants always starting with right-side cues, and others with left-side cues. Each block comprised 120 trials, with 30 trials per run. A total of 480 trials were collected from each participant.

### 2.3. Behavioural Analysis

To examine the quality of behavioural data, *response error magnitude* in each trial was computed as the angular difference between the participant’s response and the objectively correct location or orientation of the cued item. The error magnitude for the orientation task ranged between 0° and ±90°. In the location task, by contrast, the error magnitude ranged between 0° and ±180°, with errors larger than ±90° indicating that participants selected a location in the uncued hemifield. Initial data inspection indicated that participants never made such “hemifield-swap” errors, so subsequent analyses considered location in the same manner as orientation, with error magnitudes ranging between 0° and ±90°. For both tasks, the error magnitudes were transformed from degrees to pi radians (πrad) with 0° and ±90° mapped to 0 πrad and ±1 πrad, respectively. The error distribution for each participant was then compared against a uniform distribution using the Kolmogorov-Smirnov test (Massey, 1951). A uniform distribution is expected if a participant guesses in a majority of experimental trials. Based on this criterion, eight participants were excluded from further analyses, leaving a total of 72 participants (36 females; 18-38 years; *M* = 24.31, *SD* = 4.77) for the following analyses.

To quantify performance on the task, a probabilistic model introduced by Bays et al. (2009) was applied to trials where participants made a response. This model attributes response error to a mixture of three components: (a) the von Mises distribution for the target orientation/location, (b) the von Mises distribution for the non-target orientations/locations, and (c) a uniform distribution for random guesses. Technical details of this model have been described elsewhere (Bays et al., 2009). In brief, the model is defined as the probability of reporting the target item (P_*T*_), the probability of reporting the non-target items (P_*NT*_), the probability of random guessing (P_*G*_), and the concentration parameter *κ* of the von Mises distribution that described the variability around the target value. The non-target items in both the location and orientation tasks were defined as the two unprobed items presented on the cued side. The maximum likelihood estimates of the parameters were obtained separately for each participant in each task and cued side using an expectation-maximization algorithm. The fitted von Mises *κ* was converted to circular standard deviation (σ_vM_) as defined by Fisher (1995), giving a measure of *response precision* which reflects the precision of representations stored in visual working memory (Bays, Gorgoraptis, et al., 2011; Bays, Wu, et al., 2011; Pratte et al., 2017). P_*G*_ measures the *random guess rates* which reflect the proportion of responses originating from task-irrelevant factors.

A preliminary repeated-measures ANOVA on response precision, with task (location, orientation) and cued side (left, right) as the within-subject variables, showed no significant main effect of cued side and no significant interaction between task and cued side (all *ps* >> 0.05). Therefore, the mixture model was refitted to error distributions with trials aggregated across cued sides. Paired-samples *t*-tests were then performed to compare σ_vM_ and P_*G*_ between the location and orientation tasks. The task effect on P_*NT*_, or so-called “*swap errors*” describing a feature binding anomaly in working memory where a non-target feature is “swapped in” for the target feature (Bays, 2016; Schneegans & Bays, 2017) was also investigated. The mixture distribution modelling was performed in MATLAB R2020a using the Analogue Report Toolbox (Bays et al., 2009; Schneegans & Bays, 2016). All statistical analyses were carried out using R v4.1.0 (R Core Team, 2021).

### 2.4. Neuroimaging Analysis

#### 2.4.1. Image Acquisition

Participants underwent MRI scans using a Siemens Magnetom Prisma 3T system at the Centre for Advanced Imaging at The University of Queensland. T1-weighted structural scans were obtained with a magnetisation-prepared two rapid acquisition gradient echo (MP2RAGE) sequence (Marques et al., 2010), with 240 mm field-of-view (FoV), 176 slices, 0.9 mm isotropic resolution, TR = 4000 ms, TE = 2.92 ms, TI 1 = 700 ms, TI 2 = 2220 ms, first flip angle = 6°, second flip angle = 7°, and 5-6 minutes of acquisition time. Diffusion-weighted image (DWI) series were acquired using an echo-planar imaging (EPI) sequence with FoV of 244 mm, 70 slices, 2 mm isotropic resolution, TR = 4100 ms, TE = 84 ms, in 106 diffusion-sensitization directions. Diffusion weightings of b = 0, 200, 500, 1000, and 3000 s/mm^2^ were applied in 10, 6, 10, 20, and 60 directions, respectively. In addition, 11 b = 0 s/mm^2^ images were acquired with reverse phase encoding. The DWI sequence lasted approximately 7-8 minutes.

#### 2.4.2. Image Pre-processing

Image pre-processing was conducted using tools in MRtrix3 (v3.0_RC3; Tournier et al., 2019) and FSL (v6.0.4 FMRIB Software Library; Smith et al., 2004). DWI were denoised and corrected for susceptibility induced field, subject movement, eddy-current induced distortions, and signal intensity inhomogeneities (Andersson & Sotiropoulos, 2016; Cordero-Grande et al., 2019; Veraart et al., 2016; Zhang et al., 2001). A binary whole-brain mask was generated from the pre-processed DWI using the Brain Extraction Tool (BET; Smith, 2002). T1-weighted images were brain extracted with the HD-BET algorithm, a new tool that has been validated on several large datasets and multiple MR sequences (Isensee & Hucho, 2019). Compared with similar tools, HD-BET provided a better automated extraction of our T1 images. Manual editing of the extracted brain mask was applied when necessary. Finally, the DWI were registered to the MNI 152 standard space with the B_0_-to-T1 and T1-to-standard transforms (Jenkinson et al., 2002).

#### 2.4.3. Probabilistic Tractography

The SLF I, SLF II, SLF III, IFOF, and ILF in both hemispheres were reconstructed according to a standardised protocol using probabilistic tractography as implemented in the XTRACT toolbox in FSL (Fig. 2; de Groot et al., 2013; Warrington et al., 2020). To guide tractography, a ball-and-stick, crossing-fibre model was applied to the pre-processed DWI to estimate multiple fibre orientations in each voxel (Behrens et al., 2007; Jbabdi et al., 2012). Model parameters were estimated using a Bayesian Monte Carlo sampling technique. Probabilistic fibre tracking was then achieved by drawing sample streamlines from a seed along a diffusion orientation sampled from the posterior distribution at each voxel. A large number of samples built up a fibre probability distribution that reflected the number of streamlines connecting any single voxel to the seed masks (Behrens et al., 2007; Behrens et al., 2003). All parameters that constrained streamline propagation were set as default: curvature threshold = ±80°, maximum streamline steps = 2000, step size = 0.5 mm (Warrington et al., 2020). To minimize the impact of partial volume contamination, diffusion tensor images corrected for cerebrospinal fluid (CSF) were obtained with a bi-tensor model as described in Pasternak et al. (2009). The resultant CSF-corrected images were used to generate diffusion tensor maps, from which the estimated mean of *fractional anisotropy* (FA), *mean diffusivity* (MD), *radial diffusivity* (RD), and *axial diffusivity* (AD) were extracted for all TOIs.

**Figure 2.**
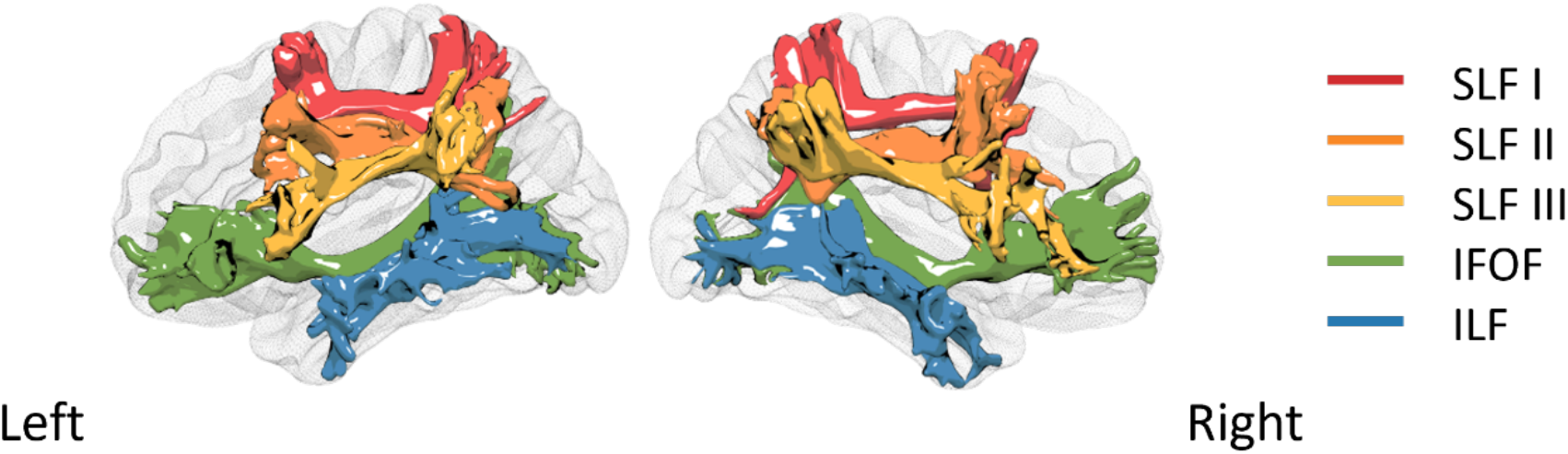
Probabilistic Tractography of Five Tracts of Interest in Both Hemispheres. Normalised streamline density maps from a representative participant were converted to 3D meshes for the purpose of visualization. *Abbreviations:* SLF, superior longitudinal fasciculus; IFOF, inferior frontal-occipital fasciculus; ILF, inferior longitudinal fasciculus.

#### 2.4.4. Principal Component Analysis on Tractography Data

The tract-specific microstructural measures derived from each participant yielded a total of 40 variables (4 diffusion tensor measures × 5 tracts × 2 hemispheres). To decrease dimensionality of the microstructure data, PCA was applied to the tractography dataset. To identify principal components (PCs) that reflected features in both the measure and tract spaces, PCA was performed in a two-step fashion (Fig. 3). The first PCA was performed on the concatenated data with participants and white matter tracts as observations (rows) and the four diffusion tensor metrics as features (columns). The second PCA was performed on each of the PCs extracted from the first PCA with the ten white matter tracts as features and participants as observations. Two statistical tests were performed before each PCA to determine whether the data were suitable for structure detection. The Kaiser-Meyer-Olkin (KMO) Statistic measures the proportion of variance that might be caused by underlying factors. High KMO values generally suggest the suitability of applying PCA (Kaiser & Rice, 1974). A KMO value lower than 0.5 is considered to render PCA inappropriate (Dziuban & Shirkey, 1974). Bartlett’s Test of Sphericity determines whether variables in a dataset are correlated; if all variables were unrelated, a PCA would be inappropriate (Bartlett, 1951). The above analyses were conducted using the ‘REdaS’ package (Maier, 2015). To obtain PCA scores and loadings, singular value decomposition was applied to the mean-centred, *z*-transformed data matrix using the ‘mdatools’ package (Kucheryayskiy, 2020). PCs with eigenvalue > 1 were retained (Cattell, 1966). Individual variables were considered to contribute substantially to a PC if they explained more variance than that would be equally explained by each variable (PC loading > 1/number of variables).

**Figure 3.**
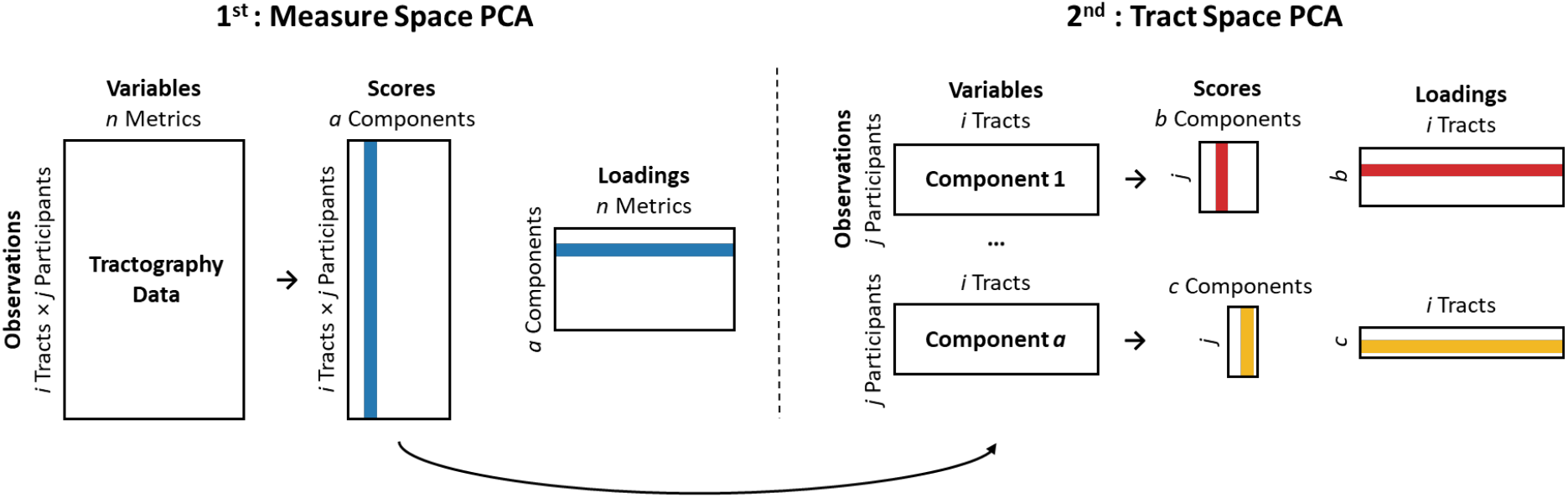
Schematic Illustration of Two-step Principal Component Analysis (PCA). The measure space PCA aimed to capture covariations across different diffusion metrics with participant and tract data as observations. Scores of the extracted components were then re-structured for the tract space PCA which captured covariations across white matter tracts.

### 2.5. Statistical Analysis

To examine brain-behaviour relationships, linear mixed effects models were used to separately regress response precision (σ_vM_) and random guess rates (P_*G*_) on the identified PCs using R packages ‘lme4’ (Bates et al., 2015) and ‘lmerTest’ (Kuznetsova et al., 2017). The white matter underpinning of swap errors (P_*NT*_) was also investigated. To find the most parsimonious model that provided the best fit to the data, we adopted a step-up model building approach. This procedure starts with the construction of a base model, followed by the stepwise addition of predictor variables. Every new model is evaluated against a simpler model via the likelihood ratio test (LRT). In the present study, we tested fixed main effects of task (location vs. orientation) and all extracted PCs characterising white matter connectivity as well as their interactions. In the case of significant interactions, the corresponding main effect terms were also retained, even if the main effects were not statistically significant. Significant interactions were followed up using simple slope analysis as implemented in the ‘reghelper’ package (Hughes, 2021). Significance level was set at *p* < 0.05; *p*-values for fixed effects were calculated using Satterthwaite approximations.

## 3. Results

### 3.1. Behavioural Results

Distributions of error magnitudes were unimodal and centred on zero for both tasks (Fig. 4), indicating that participants were successful in reporting features of the target item. On average across participants, the estimated response precision was higher for the location task than for the orientation task (σ_vM_ = 0.40/0.01 [M/SEM] vs. 0.74/0.03, *t*(71) = −10.40, *p* < 0.001). The estimated random guess rates were lower for the location task than the orientation task (P_*G*_ = 0.02/ 0.01 vs. 0.36/0.03, *t*(71) = 13.20, *p* < 0.001). Finally, the estimated swap errors were higher for the location task than the orientation task (P_*NT*_ = 0.34/0.01 vs. 0.03/0.01, *t*(71) = −28.10, *p* < 0.001).

**Figure 4.**
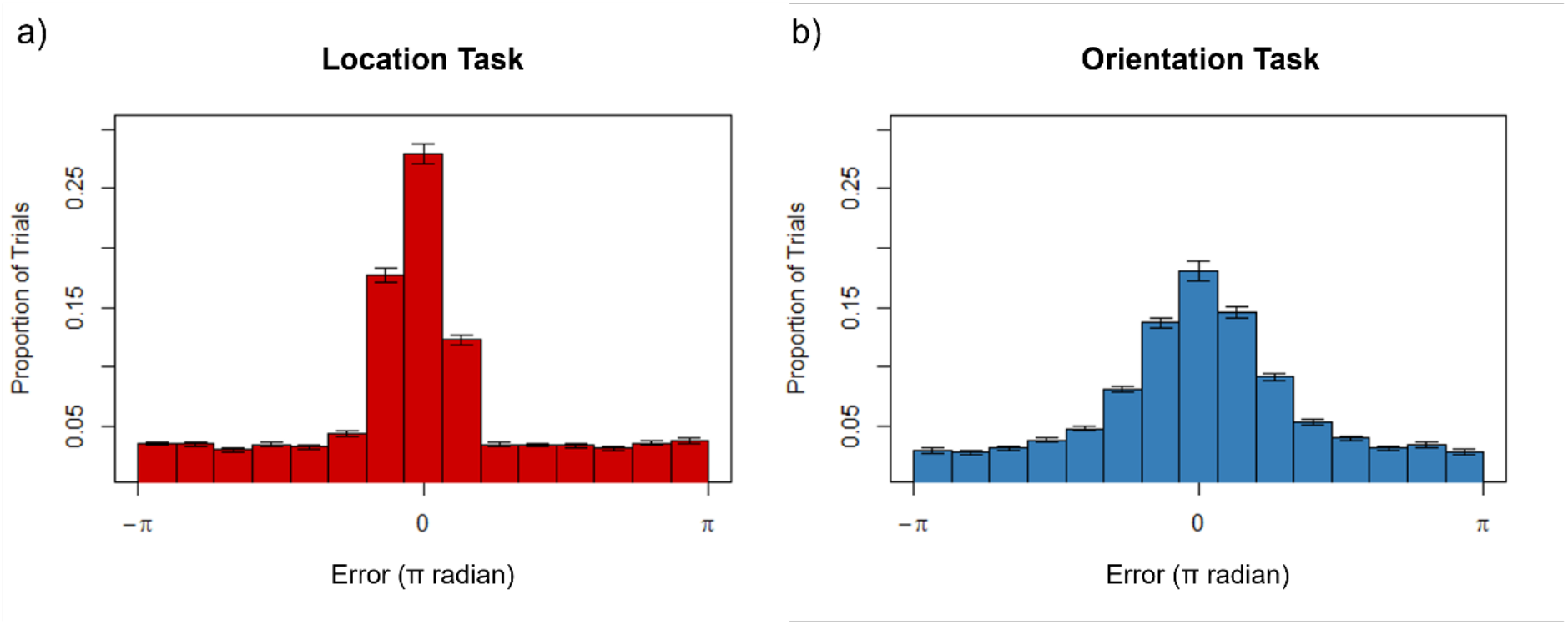
Distributions of Response Errors in Visual Working Memory Tasks. Results of the a) location task and b) orientation task are shown. Proportion of trials within each bin is averaged across participants. Error bars in each bin denote standard errors of the mean.

### 3.2. Principal Component Analysis on Tractography Data

The PCA over diffusion tensor measures identified two PCs that collectively accounted for 99.73% of the variance (KMO: 0.50; Bartlett’s Test: χ^2^ = 8389.99, *p* < 0.001). The first component (PC_1_) explained 74.5% of the variance, with FA and AD loading negatively and RD loading positively (Table S1). As higher scores on PC_1_ reflect a lower extent of anisotropic diffusion (Basser, 1995; Beaulieu, 2002), PC_1_ therefore represents decreases in *directionality*. The loading profile of PC_1_ was replicated by a complementary PCA on the three eigenvalues of the diffusion tensor (see Supplementary Results). The second PC (PC2) explained 25.2% of the variance, with MD loading negatively (Table S1). PC2 was therefore considered to represent the bulk mean magnitude of the diffusion process (labelled as “*bulk diffusion*”). To facilitate subsequent analysis, scores of PC2 were multiplied by −1 to represent increases in bulk diffusion. In brief, the measure-space PCA yielded a two-component solution, each of which represents a distinct aspect of the diffusion process.

Participant scores for each of the extracted PCs from the measure-space PCA were next submitted to the tract-space PCA to capture covariance across TOIs. For directionality, the tract-space PCA extracted three orthogonal PCs (eigenvalues > 1) which collectively explained 63.1% of the variance (KMO: 0.69; Bartlett’s Test: χ^2^ = 219.17, *p* < 0.001), and loaded onto three distinct groups of tracts (Fig. 5; Table S1). For bulk diffusion, the tract-space PCA yielded a clear single component that explained 71.5% of the variance (KMO: 0.92; Bartlett’s Test: χ^2^ = 726.50, *p* < 0.001), with all TOIs loading equally and positively on the extracted component (Fig. 5; Table S1). To summarize, four structural components, namely PC_*DIR*,1_, PC_*DIR*,2_, PC_*DIR*,3_, and PC_*BULK*_, were extracted from the tractography data.

**Figure 5.**
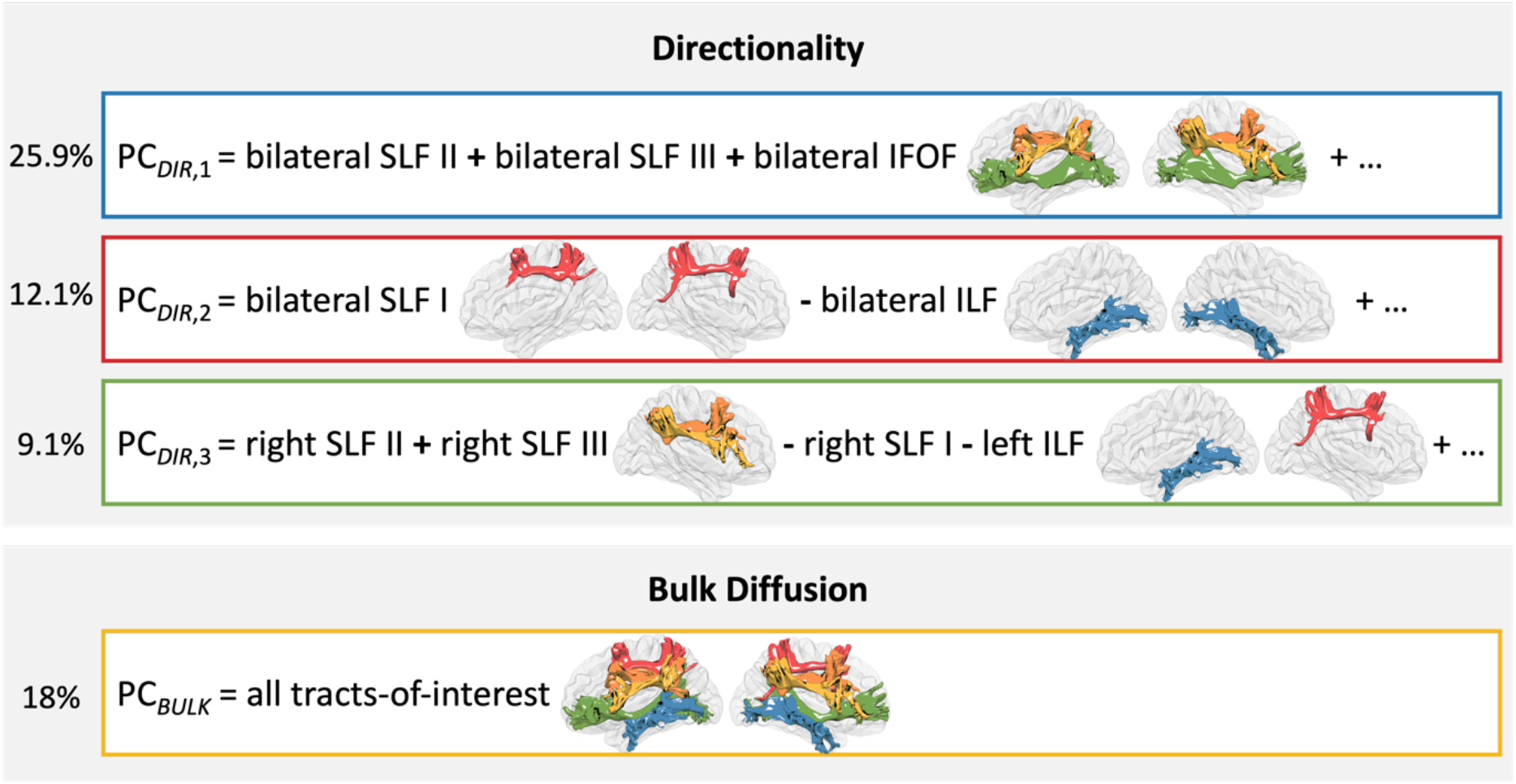
Loading Profile of the Four Extracted Components. Percentages on the left denote the proportion of variance in the whole tractography dataset explained by each principal component. *Abbreviations*: SLF, superior longitudinal fasciculus; IFOF, inferior frontal-occipital fasciculus; ILF, inferior longitudinal fasciculus.

### 3.3. Associations between Structural Components and Behavioural Performance

The model comparison and selection processes for all dependent variables are summarized in Tables S2-S4. The best-fitting model of response precision showed that PC_*BULK*_ correlated positively with σ_vM_ indicating that higher bulk diffusion across all TOIs was associated with lower response precision (Fig. 6a; Table 1). No significant interaction between task and PC_*BULK*_ was found, suggesting that the relationship between σ_vM_ and PC_*BULK*_ was similarly strong across the location and orientation tasks. In addition, there was a statistically significant interaction between PC_*DIR*,1_ and PC_*DIR*,3_ (Fig. 6b; Table 1). This interaction was followed up by using the simple slope analysis with PC_*DIR*,1_ as the main predictor and PC_*DIR*,3_ as a moderator variable. The analysis showed that lower scores in PC_*DIR*,1_ were related to lower σ_vM_ across tasks when PC_*DIR*,3_ scores were high (*p* = 0.003). When scores of PC_*DIR*,3_ were low, however, there was no statistically significant association between PC_*DIR*,1_ and σ_vM_ (*p* = 0.300). A *post hoc* model that included a three-way interaction between PC_*DIR*,1_, PC_*DIR*,3_, and task did not significantly improve the model fit, suggesting that the interaction between PC_*DIR*,1_ and PC_*DIR*,3_ did not differ between the location and orientation tasks (Tables S5-S6).

**Figure 6.**
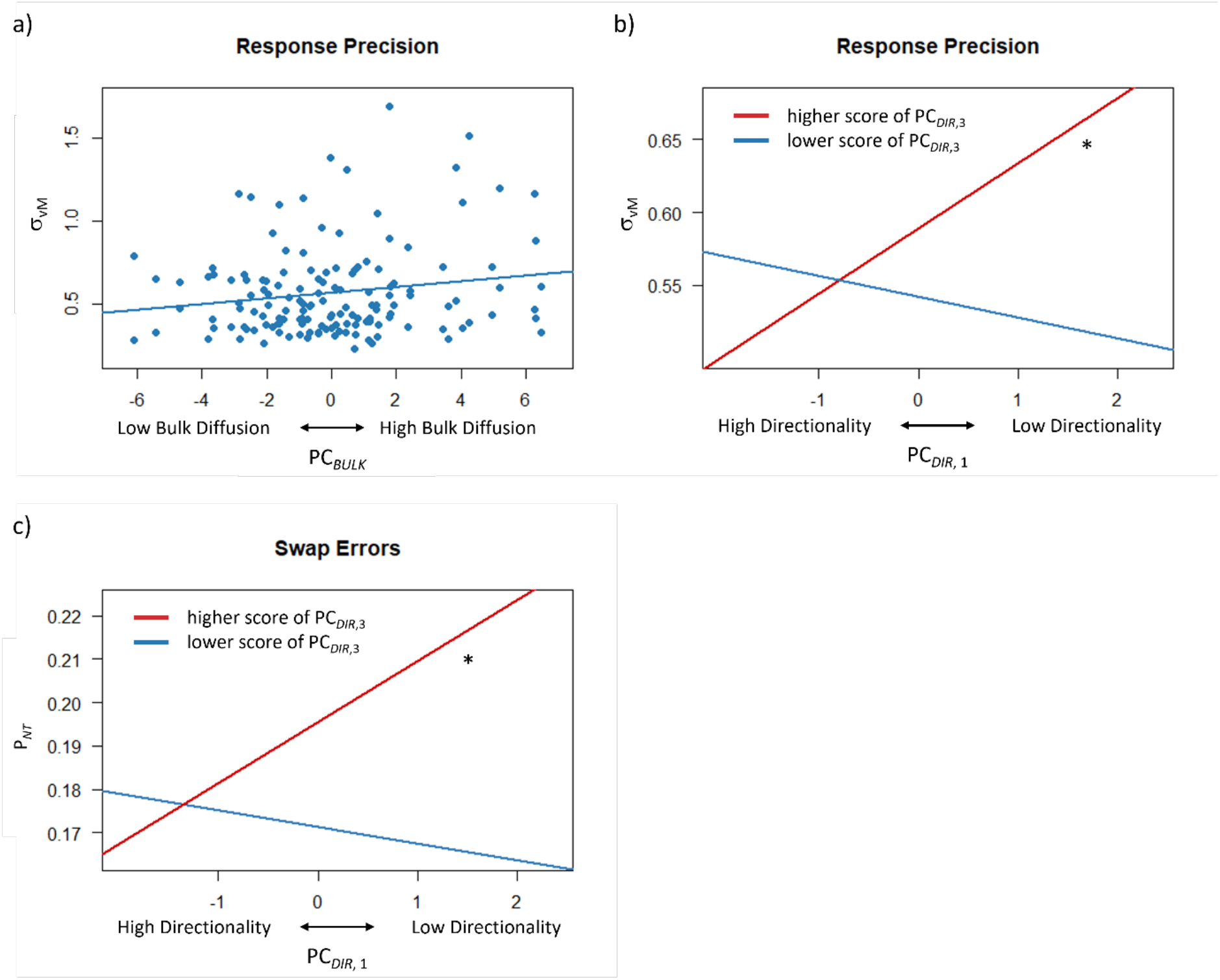
Brain-behaviour Associations Revealed by Linear Mixed Models. In the best-fitting model of response precision, a) significant effects of PC_*BULK*_ and b) significant interaction between PC_*DIR*,1_ and PC_*DIR*,3_ were found. Note that higher values in σ_vM_ represent lower response precision. Higher values in PC_*DIR*,1_ and PC_*DIR*,3_ relate to lower extent of diffusion directionality. In the best-fitting model of swap errors, c) significant interaction between PC_*DIR*,1_ and PC_*DIR*,3_ was found. Asterisks in plots b) and c) denote significant simple effects.

**Table 1.** The Best-fitting Model of Response Precision.

| Fixed Effects | Estimate | SE | *t* | 95% CI | *p* |
| --- | --- | --- | --- | --- | --- |
| Intercept (Orientation) | -0.74 | 0.02 | 30.49 | [0.69, 0.78] | < 0.001 |
| Task (Location) | -0.34 | 0.03 | -10.41 | [-0.41, -0.28] | < 0.001 |
| $PC_{BULK}$ | 0.02 | 0.01 | 2.25 | [0.00, 0.03] | 0.028 |
| $PC_{DIR,1}$ | 0.02 | 0.01 | 1.50 | [0.00, 0.03] | 0.138 |
| $PC_{DIR,3}$ | 0.02 | 0.02 | 1.30 | [-0.01, 0.05] | 0.200 |
| $PC_{DIR,1} \times PC_{DIR,3}$ | 0.03 | 0.01 | 3.02 | [0.01, 0.04] | 0.004 |

The best-fitting model of random guess rates, by contrast, did not show any significant effect of the structural components (all *ps* >> 0.05), suggesting that the brain-behaviour associations we observed are functionally specific to the precision of visual working memory independently of random guessing.

The best-fitting model of swap errors revealed a statistically significant interaction between PC_*DIR*,1_ and PC_*DIR*,3_ (Fig. 6c; Table 2). The interaction was followed up in the same way as in the model of response precision to investigate the relationship between PC_*DIR*,1_ and swap errors at high and low levels of PC_*DIR*,3_. The analysis showed that lower scores in PC_*DIR*,1_ were related to lower probability of swap errors when scores of PC_*DIR*,3_ were higher (*p* = 0.011). When scores of PC_*DIR*,3_ were lower, there was no association between PC_*DIR*,1_ and swap errors (*p* = 0.440). A *post hoc* model that included a three-way interaction between PC_*DIR*,1_, PC_*DIR*,3_, and task did not significantly improve the model fit (Tables S7-S8).

**Table 2.**
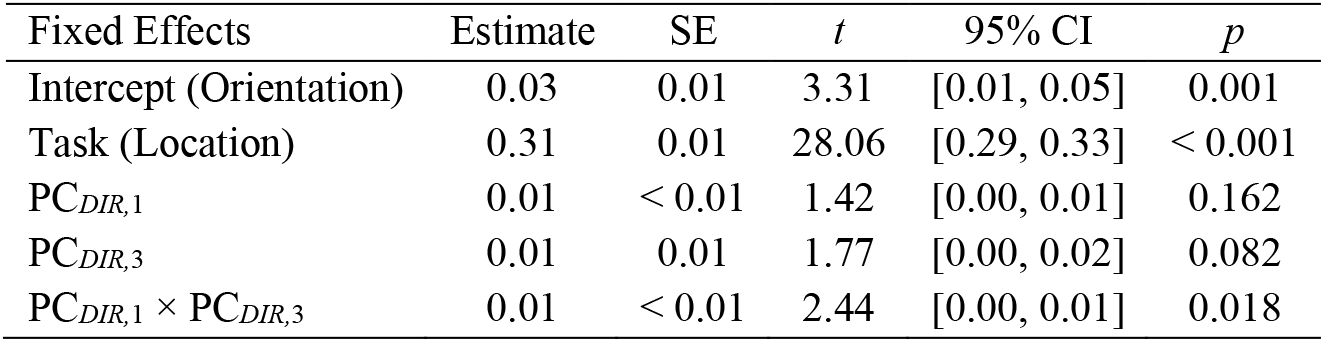
The Best-fitting Model of Swap Errors.

## 4. Discussion

The principal aim of the current study was to characterise the relationship between distinct aspects of visual working memory performance and the microstructure of major white matter tracts in the healthy adult brain. We found that components of tract-specific microstructural properties were associated with the precision of visual working memory but not with occasional random guesses made by participants during the tasks. Higher response precision was associated with lower bulk diffusion shared across all reconstructed tracts. Higher response precision, as well as lower swap errors, were associated with higher directionality in a set of bilateral frontoparietal-occipital tracts in individuals with decreased directionality in particular right frontoparietal tracts.

In the best-fitting model of response precision, we found that lower scores on PC_*BULK*_, most likely reflecting lower bulk diffusion across white matter tracts, were associated with higher precision in both the location and orientation tasks. It is noteworthy that the models of swap errors and random guess rates, by contrast, showed no effect of PC_*BULK*_, suggesting that the general bulk diffusion of white matter tracts specifically mediates the precision of visual working memory and not task-irrelevant factors such as attention lapses. A previous study in children has shown that higher capacity in visual working memory task was associated with decreased MD in the IFOF and ILF (Krogsrud et al., 2018). A recent study in older adults identified a negative association between global MD of whole-brain white matter and the sensitivity index in visual working memory task (Conley et al., 2020). Our findings go beyond this work by showing a specific relationship between general bulk diffusion and precision of memory representations rather than merely the overall capacity or accuracy of working memory. This finding also suggests that the precision of working memory representations may rely on a general, trait-like property of white matter tracts that is reflected by individual differences in bulk diffusion. Axon density, for example, is one possible candidate. This property, being negatively related to MD, determines how much information can be carried by the white matter tract (Alexander et al., 2007; Alexander et al., 2019; Beaulieu, 2002). Increases in axon density in young adulthood across all major pathways and global white matter are much slower than those in childhood and adolescence (Chang et al., 2015). Axon density may therefore serve as a relatively static, trait-like property in our sample by imposing resource limitations on visual working memory. In this case, our findings suggest that people with higher axonal density have more precise responses when reproducing features from visual working memory. It should be noted, however, that the measure of bulk diffusion/MD is also sensitive to other microstructural attributes such as axonal diameter and membrane permeability (Jones et al., 2013). Future research could extend our work using more sophisticated DWI models such as the neurite orientation dispersion and density imaging (NODDI) model (Zhang et al., 2012).

The model of response precision also showed a significant interaction between PC_*DIR*,1_ and PC_*DIR*,3_. Higher scores on these components describe a lower extent of directional diffusion along the axonal fibres, which is generally found in white matter tracts that are less myelinated or less coherently packed (Alexander et al., 2007; Beaulieu, 2002; Jones et al., 2013). Critically, both components were related to some but not all TOIs in our study, which suggests the presence of shared microstructural properties in specific groups of tracts. PC_*DIR*,1_ represents directionality in a set of bilateral frontoparietal-occipital tracts (i.e., SLF II, SLF III, and IFOF in both hemispheres). Two of these tracts (i.e., the right SLF II and right SLF III) also contributed substantially to PC_*DIR*,3_. We found that higher directionality in the frontoparietal-occipital tracts was associated with higher memory precision in both location and orientation tasks in individuals with decreased directionality in the right SLF II and right SLF III. Previous studies have shown that increased FA in the SLF and IFOF was associated with better performance in visual working memory tasks (Darki & Klingberg, 2015; Peters et al., 2014; Vestergaard et al., 2011; Walsh et al., 2011). Our result further extends the previous findings by showing a specific relationship between diffusion directionality and the precision of visual working memory, independently of occasional random guesses originating from task-irrelevant factors. The interaction effect between PC_*DIR*,1_ and PC_*DIR*,3_ also reveals the interdependence of different sets of white matter tracts when predicting response precision in visual working memory tasks. This observation suggests that individual differences in working memory precision are modulated by the complex interplay between subsets of tracts across a wider working memory network.

The interaction between PC_*DIR*,1_ and PC_*DIR*,3_ was also associated with swap errors. Higher directionality in the frontoparietal-occipital tracts predicted lower swap errors across tasks in individuals with relatively low directionality in the right SLF II/III. The estimate of swap errors reflects a specific anomaly in feature binding between the probed and reported features (Schneegans & Bays, 2017). Previous studies of visual working memory have implicated the superior parietal area, inferior intraparietal sulcus, and lateral occipital regions in feature binding (Parra et al., 2014; Shafritz et al., 2002; Xu & Chun, 2006). White matter microstructure in the inferior frontal cortex has also been associated with feature binding performance (Parra et al., 2015). Binding deficits have been selectively associated with lesions in the left somatosensory cortex (Lugtmeijer et al., 2021). Our results further extend previous work by showing that binding errors that occur during visual working memory tasks are modulated by the microstructure of long-range white matter tracts that connect the frontal, parietal, and occipital regions.

The fact that we observed a significant interaction between PC_*DIR*,1_ and PC_*DIR*,3_ when modelling both response precision and swap errors implies common neural substrates for these distinct components of visual working memory performance. A recent study suggests that both response variability and swap errors are caused by stochastic noise in neural activity (Schneegans & Bays, 2017), for example, variability in neural spike timing (Faisal et al., 2008). Although the presence of noise is universal in the nervous system, some individuals may benefit from a relatively high signal-to-noise ratio, which can be modulated by, for example, the level of axonal myelination. When compared with unmyelinated axons, those insulated by myelin sheaths can modulate the speed of impulse conduction and thereby facilitate optimal synchronization among neural assemblies in distant regions (Fields & Bukalo, 2020; Nunez et al., 2015; Pajevic et al., 2014). Individuals with more myelinated axons in general may have more fine-grained modulation on conduction speed, which in turn allows a wider spectrum of synchronized oscillations. Since long-range, inter-areal synchronization during visual working memory tasks has been found for a wide range of frequency bands (Daume et al., 2017; Palva et al., 2010; Sato et al., 2018), greater myelination might therefore contribute to more precise responses and lower swap errors by boosting signal resolution.

One other interesting observation of the interaction effects is that people with lower scores on PC_*DIR*,3_ tended to have overall higher response precision and lower swap errors (Fig. 6b-c, blue lines). Such good performance was not related to scores on PC_*DIR*,1_, which suggests that higher directionality along the right SLF II and SLF III operates as a protective factor, shielding visual working memory from effects arising in the set of bilateral frontoparietal-occipital tracts. When such protection is weak, that is, when microstructure of the right SLF II/III is somehow suboptimal, individual differences in task performance were found to covary strongly with differences in directionality in the bilateral frontoparietal-occipital network. The strong reliance on long-range white matter tracts across a large bilateral network may impose a higher degree of susceptibility to disturbance in visual working memory perhaps due to focal lesions in white matter arising from neurological diseases. To speculate more broadly, our findings also suggest that swap errors occur, at least in a significant portion of trials, when the precision of the maintained representation is low, which in turn increases confusability between different features stored in visual working memory. This claim is consistent with previous finding that elevated swap errors were observed when the target item was more similar to the non-target distractors (Schneegans & Bays, 2017).

The observed effect of PC_*BULK*_ and the interaction between PC_*DIR*,1_ and PC_*DIR*,3_ did not show any notable differences between the location and orientation tasks, even though we observed robust differences between tasks when we analysed behavioural data only. In addition, our study comprised a relatively large sample, suggesting that we should have been able to detect even small differences between tasks in brain-behaviour relationship should these differences exist. The location and orientation tasks in our study involved a common encoding display but in which participants were instructed to reproduce either the spatial or non-spatial features. Previous studies using EEG recording have shown that the contralateral delay activity (CDA) tracks different features stored in visual working memory. When participants were presented with an identical array of items, greater CDA was found when participants were tested on the orientations or shapes of the items, compared with when they were tested on the colours of the items (Luria et al., 2010; Woodman & Vogel, 2008). This feature-specific difference, however, was not apparent in the microstructural properties in long-range white matter tracts in our study. Recent studies have argued that human cognition arises from dynamic interactions within and between large-scale networks rather than coming from several discrete, specialised brain regions (Bassett & Sporns, 2017; Bressler & Menon, 2010; Park & Friston, 2013). Anatomical connections between different brain areas, comprising relatively invariant white matter fibres, predict but do not fully determine the dynamic repertoire of cognitive functions (Honey et al., 2009; Suarez et al., 2020). The functional differences between the location and orientation tasks may therefore arise from neural mechanisms that are not constrained by or sensitive to white matter microstructure.

In this study, we applied a two-step PCA to the tractography data which extracted four orthogonal PCs that encompassed critical information in both the measure and tract spaces. This approach addressed problems that were overlooked in previous studies investigating the relationship between visual working memory and white matter tracts. For example, Krogsrud and colleagues (2018) failed to find associations between visual working memory performance and the tract-based FA measures after correcting for multiple comparisons. The fact that Krogsrud et al.’s study did not replicate otherwise reproducible findings (e.g., Darki & Klingberg, 2015; Nagy et al., 2004; Peters et al., 2014; Vestergaard et al., 2011; Walsh et al., 2011), raises an issue of high dimensionality in the predictor space. The use of PCA in our study helped achieve data reduction while maintaining maximal information in the tractography dataset. In addition, although all studies have made claims that certain tracts are related to visual working memory, few studies have controlled for the general effect of individual differences in white matter microstructure (Krogsrud et al., 2018; Vestergaard et al., 2011; Walsh et al., 2011), leading to inconclusive results regarding tract specificity. The “global” microstructure can explain up to 73% of the variance and therefore become non-negligible in some datasets (Clayden et al., 2012; Johnson et al., 2015; Penke et al., 2010). The application of tract-space PCA effectively identified the global microstructural component (PC_*BULK*_) in our dataset, which helped find effects that are above and beyond the general differences in bulk diffusion. To summarize, the two-step PCA approach in our study highlights the fact that the modulating influence of white matter microstructure goes beyond the level of individual metrics or tracts.

## 5. Conclusions

In the present study, we observed brain-behaviour associations between visual working memory and white matter microstructure that were functionally specific to the precision of representations in visual working memory, both spatial and non-spatial. Higher response precision was associated with lower bulk diffusion across all reconstructed white matter tracts. Response precision was also positively related to diffusion directionality in a particular group of bilateral frontoparietal-occipital pathways, but these associations depended on microstructural properties of another group of tracts. Our findings suggest that individual differences in the precision of visual working memory reflect inter-subject variability in both widespread and regional properties of white matter microstructure, which in turn show evidence of interaction effects across the wider working memory network.

## Acknowledgements

The authors would like to thank Dr David Lloyd from the University of Queensland for his assistance with the experimental set-up.

## CRediT Author Statement

**Xuqian Li:** Conceptualization, Methodology, Software, Investigation, Writing - Original Draft, Writing - Review & Editing, Visualization; **Dragan Rangelov:** Conceptualization, Methodology, Software, Writing - Review & Editing; Supervision; **Jason B. Mattingley:** Conceptualization, Methodology, Writing - Review & Editing, Supervision; **Lena Oestreich:** Methodology, Writing - Review & Editing; **Delphine Lévy-Bencheton:** Conceptualization, Methodology, Investigation; **Michael J. O’Sullivan:** Conceptualization, Methodology, Writing - Review & Editing, Supervision.

## Funding Sources

XL was supported by a scholarship from the Graduate School for Research Training, The University of Queensland. DR was supported by a National Health and Medical Research Council (Australia) Ideas grant (APP1186955). MOS was supported by a strategic award from the Deputy Vice-Chancellor for Research and Innovation, The University of Queensland. JBM was supported by a National Health and Medical Research Council (Australia) Investigator Grant (GNT2010141).

## Data Availability Statement

Data used in this study are available for download from the OSF repository (https://osf.io/4wkuf/?view_only=9ccba5df6110466590077ea615614f4f).

## Conflict of Interest

The authors declare that there is no conflict of interests.

## Notes

https://osf.io/4wkuf/?view_only=9ccba5df6110466590077ea615614f4f

### Competing Interest Statement

The authors have declared no competing interest.

