## Supplement for "White matter microstructure is associated with the precision of visual working memory"

**Supplementary Materials**

**Supplementary Results.** Complementary PCA on Diffusion Tensor Eigenvalues.

**Table S1.** PCA Loadings of the Measure Space PCA and Tract Space PCA.

**Table S2.** Model Comparison and Selection: Response Precision.

**Table S3.** Model Comparison and Selection: Swap Errors.

**Table S4.** Model Comparison and Selection: Random Guesses.

**Table S5.** The Post-hoc Model of Response Precision.

**Table S6.** Model Comparison: The Best-fitting Model *vs.* Post-hoc Model of Response Precision.

**Table S7.** The Post-hoc Model of Swap Errors.

**Table S8.** Model Comparison: The Best-fitting Model *vs.* Post-hoc Model of Swap Errors.

**Supplementary Results.** Complementary PCA on Diffusion Tensor Eigenvalues.

The loading profile of measure space PC_1_ was replicated by a complementary PCA on the three eigenvalues of the diffusion tensor (KMO: 0.66; Bartlett’s Test: χ^2^ = 2212.05, *p* < 0.001). The analysis extracted one PC that explained 89.2% of the variance. Loadings on this PC suggested a negative association with the major eigenvector (loading = -0.598) and positive associations with the second and third eigenvectors (loadings = 0.563 and 0.57, respectively), consistent with the loading pattern of PC_1_.

**Table S1.** PCA Loadings of the Measure Space PCA and Tract Space PCA.

| Measure Space PCA | | |  | Tract Space PCA | | | | |
| --- | --- | --- | --- | --- | --- | --- | --- | --- |
|  | PC_1_ | PC_2_ |  |  | PC*_DIR,_*_1_ | PC*_DIR,_*_2_ | PC*_DIR,_*_3_ | PC*_BULK_* |
| FA | **-0.574** | 0.103 |  | IFOF L. | **0.408** | -0.307 | 0.005 | **0.323** |
| MD | -0.149 | **-0.962** |  | IFOF R. | **0.388** | -0.264 | 0.155 | **0.329** |
| RD | **0.562** | -0.237 |  | ILF L. | 0.103 | **-0.389** | **-0.509** | **0.323** |
| AD | **-0.577** | -0.084 |  | ILF R. | 0.190 | **-0.559** | -0.118 | **0.316** |
|  |  |  |  | SLF I L. | 0.284 | **0.402** | -0.248 | **0.311** |
|  |  |  |  | SLF I R. | 0.223 | **0.357** | **-0.583** | **0.309** |
|  |  |  |  | SLF II L. | **0.354** | 0.201 | 0.059 | **0.293** |
|  |  |  |  | SLF II R. | **0.336** | 0.186 | **0.389** | **0.308** |
|  |  |  |  | SLF III L. | **0.359** | 0.079 | -0.181 | **0.325** |
|  |  |  |  | SLF III R. | **0.372** | 0.048 | **0.338** | **0.323** |

*Notes*: For the measure space PCA, variable with loading > |0.5| (loading^2^ > 25%) is considered to contribute substantially. For the tract space PCA, variable with loading > |0.32| (loading^2^ > 10%) is considered to have substantial contribution. Note that variable loadings on PC*_BULK_* are very close to each other, and we considered an equal loading profile for this component. Variables showed substantial contribution are highlighted in bold.

**Table S2.** Model Comparison and Selection: Response Precision.

| Iteration | Model | Nested Model | Effect Added | REML Fit | |  | ML Fit | | | | |  | LRT Against Nested | | |
| --- | --- | --- | --- | --- | --- | --- | --- | --- | --- | --- | --- | --- | --- | --- | --- |
|  |  |  |  | LL | Order |  | df | AIC | BIC | LL | Deviance |  | χ^2^ | df | *p* |
| 1 | *Base* | - | - | 13.15 | - |  | 4 | -29.36 | -17.48 | 18.68 | -37.36 |  | - | - | - |
|  | 1 | *Base* | PC*_BULK_* | 12.05 | 1 |  | 5 | -33.32 | -18.47 | 21.66 | -43.32 |  | 5.96 | 1 | 0.015 |
| 2 | 2 | *Base +* PC*_BULK_* | PC*_BULK_* × Task | 10.06 | 1 |  | 6 | -34.37 | -16.55 | 23.18 | -46.37 |  | 3.05 | 1 | 0.081 |
|  | 3 |  | PC*_DIR,_*_1_ | 9.20 | 2 |  | 6 | -32.90 | -15.08 | 22.45 | -44.90 |  | 1.58 | 1 | 0.210 |
|  | 4 |  | PC*_DIR,_*_3_ | 9.14 | 3 |  | 6 | -31.79 | -13.98 | 21.90 | -43.79 |  | 0.47 | 1 | 0.491 |
|  | 5 |  | PC*_DIR,_*_2_ | 9.09 | 4 |  | 6 | -31.99 | -14.18 | 22.00 | -43.99 |  | 0.67 | 1 | 0.412 |
|  | 6 |  | PC*_DIR,_*_1_ + PC*_DIR,_*_1_ × Task | 7.67 | 5 |  | 7 | -34.17 | -13.38 | 24.09 | -48.17 |  | 4.85 | 2 | 0.088 |
|  | 7 |  | PC*_DIR,_*_2_ + PC*_DIR,_*_2_ × Task | 7.45 | 6 |  | 7 | -32.23 | -11.45 | 23.12 | -46.23 |  | 2.91 | 2 | 0.233 |
|  | 8 |  | PC*_DIR,_*_3_ + PC*_DIR,_*_3_ × PC*_BULK_* | 7.34 | 7 |  | 7 | -35.13 | -14.34 | 24.57 | -49.13 |  | 5.81 | 2 | 0.055 |
|  | 9 |  | PC*_DIR,_*_1_ + PC*_DIR,_*_3_ + PC*_DIR,_*_1_ × PC*_DIR,_*_3_ | 6.78 | 8 |  | 8 | -38.51 | -14.76 | 27.26 | -54.51 |  | 11.19 | 3 | 0.011 |
| 3 | 10 | *Base* + PC*_BULK_* + PC*_DIR,_*_1_ + PC*_DIR,_*_3_ + PC*_DIR,_*_1_ × PC*_DIR,_*_3_ | - | 6.78 | 1 |  | 8 | -38.51 | -14.76 | 27.26 | -54.51 |  | 0.00 | 0 | - |
|  | 11 |  | - | 6.78 | 2 |  | 8 | -38.51 | -14.76 | 27.26 | -54.51 |  | 0.00 | 0 | - |
|  | 12 |  | PC*_DIR,_*_1_ × Task | 5.26 | 3 |  | 9 | -39.79 | -13.06 | 28.89 | -57.79 |  | 3.27 | 1 | 0.070 |
|  | 13 |  | PC*_BULK_* × Task | 4.79 | 4 |  | 9 | -39.56 | -12.83 | 28.78 | -57.56 |  | 3.05 | 1 | 0.081 |
|  | 14 |  | PC*_DIR,_*_3_ × Task | 4.23 | 5 |  | 9 | -36.59 | -9.86 | 27.30 | -54.59 |  | 0.08 | 1 | 0.781 |
|  | 15 |  | PC*_DIR,_*_2_ | 3.48 | 6 |  | 9 | -36.56 | -9.83 | 27.28 | -54.56 |  | 0.04 | 1 | 0.836 |
|  | 16 |  | PC*_DIR,_*_3_ × PC*_BULK_* | 3.40 | 7 |  | 9 | -38.53 | -11.80 | 28.27 | -56.53 |  | 2.02 | 1 | 0.155 |
|  | 17 |  | PC*_DIR,_*_1_ × PC*_BULK_* | 2.75 | 8 |  | 9 | -37.91 | -11.18 | 27.96 | -55.91 |  | 1.40 | 1 | 0.237 |
|  | 18 |  | PC*_DIR,_*_2_ + PC*_DIR,_*_2_ × Task | 1.84 | 9 |  | 10 | -36.80 | -7.10 | 28.40 | -56.80 |  | 2.28 | 2 | 0.319 |
|  | 19 |  | PC*_DIR,_*_2_ + PC*_DIR,_*_2_ × PC*_DIR,_*_3_ | 0.95 | 10 |  | 10 | -36.08 | -6.38 | 28.04 | -56.08 |  | 1.56 | 2 | 0.457 |
|  | 20 |  | PC*_DIR,_*_2_ + PC*_DIR,_*_2_ × PC*_DIR,_*_4_ | 0.27 | 11 |  | 10 | -36.53 | -6.83 | 28.27 | -56.53 |  | 2.02 | 2 | 0.365 |
|  | 21 |  | PC*_DIR,_*_2_ + PC*_DIR,_*_1_ × PC*_DIR,_*_2_ | -0.14 | 12 |  | 10 | -35.04 | -5.34 | 27.52 | -55.04 |  | 0.53 | 2 | 0.769 |

*Notes*: Total observations = 144, *N* = 72.

**Table S3.** Model Comparison and Selection: Swap Errors.

| Iteration | Model | Nested Model | Effect Added | REML Fit | |  | ML Fit | | | | |  | LRT Against Nested | | |
| --- | --- | --- | --- | --- | --- | --- | --- | --- | --- | --- | --- | --- | --- | --- | --- |
|  |  |  |  | LL | Order |  | df | AIC | BIC | LL | Deviance |  | χ^2^ | df | *p* |
| 1 | *Base* | - | - | 163.38 | - |  | 4 | -334.05 | -322.17 | 171.02 | -342.05 |  | - | - | - |
|  | 1 | *Base* | PC*_DIR,_*_3_ | 160.01 | 1 |  | 5 | -333.63 | -318.78 | 171.81 | -343.63 |  | 1.58 | 1 | 0.209 |
|  | 2 |  | PC*_DIR,_*_1_ | 159.37 | 2 |  | 5 | -333.39 | -318.54 | 171.69 | -343.39 |  | 1.34 | 1 | 0.247 |
|  | 3 |  | PC*_DIR,_*_2_ | 159.15 | 3 |  | 5 | -332.17 | -317.32 | 171.08 | -342.17 |  | 0.12 | 1 | 0.732 |
|  | 4 |  | PC*_BULK_* | 159.01 | 4 |  | 5 | -333.40 | -318.55 | 171.70 | -343.40 |  | 1.35 | 1 | 0.246 |
|  | 5 |  | PC*_DIR,_*_1_ + PC*_DIR,_*_1_ × Task | 156.69 | 5 |  | 6 | -334.51 | -316.69 | 173.25 | -346.51 |  | 4.46 | 2 | 0.108 |
|  | 6 |  | PC*_DIR,_*_3_ + PC*_DIR,_*_3_ × Task | 156.56 | 6 |  | 6 | -332.10 | -314.28 | 172.05 | -344.10 |  | 2.05 | 2 | 0.359 |
|  | 7 |  | PC*_BULK_* + PC*_BULK_* × Task | 155.49 | 7 |  | 6 | -333.53 | -315.71 | 172.77 | -345.53 |  | 3.48 | 2 | 0.175 |
|  | 8 |  | PC*_DIR,_*_2_ + PC*_DIR,_*_2_ × Task | 155.46 | 8 |  | 6 | -330.42 | -312.60 | 171.21 | -342.42 |  | 0.37 | 2 | 0.830 |
|  | 9 |  | PC*_DIR,_*_1_ + PC*_DIR,_*_3_ + PC*_DIR,_*_1_ × PC*_DIR,_*_3_ | 154.10 | 9 |  | 7 | -337.01 | -316.23 | 175.51 | -351.01 |  | 8.97 | 3 | 0.030 |
| 2 | 10 | *Base* + PC*_DIR,_*_1_ + PC*_DIR,_*_3_ + PC*_DIR,_*_1_ × PC*_DIR,_*_3_ | - | 154.10 | 1 |  | 7 | -337.01 | -316.23 | 175.51 | -351.01 |  | -316.23 | 0 | - |
|  | 11 |  | - | 154.10 | 2 |  | 7 | -337.01 | -316.23 | 175.51 | -351.01 |  | -316.23 | 0 | - |
|  | 12 |  | PC*_DIR,_*_1_ × Task | 151.42 | 3 |  | 8 | -338.14 | -314.38 | 177.07 | -354.14 |  | -314.38 | 1 | 0.077 |
|  | 13 |  | PC*_DIR,_*_3_ × Task | 150.65 | 4 |  | 8 | -335.48 | -311.73 | 175.74 | -351.48 |  | -311.73 | 1 | 0.493 |
|  | 14 |  | PC*_DIR,_*_2_ | 149.82 | 5 |  | 8 | -335.05 | -311.29 | 175.53 | -351.05 |  | -311.29 | 1 | 0.845 |
|  | 15 |  | PC*_BULK_* | 149.71 | 6 |  | 8 | -336.33 | -312.57 | 176.17 | -352.33 |  | -312.57 | 1 | 0.251 |
|  | 16 |  | PC*_BULK_* + PC*_BULK_* × Task | 146.18 | 7 |  | 9 | -336.47 | -309.74 | 177.23 | -354.47 |  | -309.74 | 2 | 0.178 |
|  | 17 |  | PC*_DIR,_*_2_ + PC*_DIR,_*_2_ × Task | 146.12 | 8 |  | 9 | -333.31 | -306.58 | 175.65 | -351.31 |  | -306.58 | 2 | 0.863 |
|  | 18 |  | PC*_DIR,_*_2_ + PC*_DIR,_*_2_ × PC*_DIR,_*_3_ | 145.66 | 9 |  | 9 | -333.13 | -306.40 | 175.57 | -351.13 |  | -306.40 | 2 | 0.943 |
|  | 19 |  | PC*_DIR,_*_2_ + PC*_DIR,_*_1_ × PC*_DIR,_*_2_ | 145.03 | 10 |  | 9 | -333.12 | -306.39 | 175.56 | -351.12 |  | -306.39 | 2 | 0.951 |
|  | 20 |  | PC*_BULK_* + PC*_DIR,_*_1_ × PC*_BULK_* | 144.71 | 11 |  | 9 | -335.75 | -309.02 | 176.88 | -353.75 |  | -309.02 | 2 | 0.254 |
|  | 21 |  | PC*_BULK_* + PC*_DIR,_*_3_ × PC*_BULK_* | 144.45 | 12 |  | 9 | -334.38 | -307.66 | 176.19 | -352.38 |  | -307.66 | 2 | 0.504 |
|  | 22 |  | PC*_DIR,_*_2_ + PC*_BULK_* + PC*_DIR,_*_2_ × PC*_BULK_* | 140.44 | 13 |  | 10 | -332.58 | -302.88 | 176.29 | -352.58 |  | -302.88 | 3 | 0.668 |

*Notes*: Total observations = 144, *N* = 72.

**Table S4.** Model Comparison and Selection: Random Guesses.

| Iteration | Model | Nested Model | Fixed Effect Added | Model Fit (REML) | |  | Model Fit (ML) | | | | |  | LRT Against Nested | | |
| --- | --- | --- | --- | --- | --- | --- | --- | --- | --- | --- | --- | --- | --- | --- | --- |
|  |  |  |  | LL | Order |  | df | AIC | BIC | LL | Deviance |  | χ^2^ | df | *p* |
| 1 | *Base* | - | - | 52.39 | - |  | 4 | -108.94 | -97.06 | 58.47 | -116.94 |  |  |  |  |
|  | 1 | *Base* | PC*_DIR,_*_2_ | 49.23 | 1 |  | 5 | -109.06 | -94.22 | 59.53 | -119.06 |  | 2.12 | 1 | 0.1453 |
|  | 2 |  | PC*_DIR,_*_1_ | 46.30 | 2 |  | 5 | -108.56 | -93.71 | 59.28 | -118.56 |  | 1.62 | 1 | 0.2036 |
|  | 3 |  | PC*_DIR,_*_3_ | 49.85 | 3 |  | 5 | -107.08 | -92.23 | 58.54 | -117.08 |  | 0.13 | 1 | 0.7147 |
|  | 4 |  | PC*_BULK_* | 47.53 | 4 |  | 5 | -106.94 | -92.09 | 58.47 | -116.94 |  | 0.00 | 1 | 0.9985 |
|  | 5 |  | PC*_DIR,_*_2_ × Task | 43.21 | 5 |  | 6 | -108.39 | -90.57 | 60.20 | -120.39 |  | 3.45 | 2 | 0.1784 |
|  | 6 |  | PC*_DIR,_*_1_ × Task | 49.03 | 6 |  | 6 | -107.42 | -89.61 | 59.71 | -119.42 |  | 2.48 | 2 | 0.289 |
|  | 7 |  | PC*_DIR,_*_3_ × Task | 46.21 | 7 |  | 6 | -105.09 | -87.28 | 58.55 | -117.09 |  | 0.15 | 2 | 0.9272 |
|  | 8 |  | PC*_BULK_* × Task | 42.04 | 8 |  | 6 | -104.99 | -87.17 | 58.50 | -116.99 |  | 0.05 | 2 | 0.9757 |
|  | 9 |  | PC*_DIR,_*_1_ + PC*_DIR,_*_2_ + PC*_DIR,_*_1_ × PC*_DIR,_*_2_ | 43.14 | 9 |  | 7 | -108.06 | -87.27 | 61.03 | -122.06 |  | 5.11 | 3 | 0.1636 |
|  | 10 |  | PC*_DIR,_*_2_ + PC*_DIR,_*_3_ + PC*_DIR,_*_2_ × PC*_DIR,_*_3_ | 48.08 | 10 |  | 7 | -105.50 | -84.72 | 59.75 | -119.50 |  | 2.56 | 3 | 0.4644 |
|  | 11 |  | PC*_DIR,_*_1_ + PC*_DIR,_*_3_ + PC*_DIR,_*_1_ × PC*_DIR,_*_3_ | 44.39 | 11 |  | 7 | -105.11 | -84.33 | 59.56 | -119.11 |  | 2.17 | 3 | 0.5374 |
|  | 12 |  | PC*_DIR,_*_3_ + PC*_BULK_* + PC*_DIR,_*_3_ × PC*_BULK_* | 40.51 | 12 |  | 7 | -106.17 | -85.38 | 60.09 | -120.17 |  | 3.23 | 3 | 0.3578 |
|  | 13 |  | PC*_DIR,_*_2_ + PC*_BULK_* + PC*_DIR,_*_2_ × PC*_BULK_* | 41.26 | 13 |  | 7 | -105.20 | -84.42 | 59.60 | -119.20 |  | 2.26 | 3 | 0.5198 |
|  | 14 |  | PC*_DIR,_*_1_ + PC*_BULK_* + PC*_DIR,_*_1_ × PC*_BULK_* | 41.61 | 14 |  | 7 | -105.49 | -84.70 | 59.74 | -119.49 |  | 2.54 | 3 | 0.4673 |

*Notes*: Total observations = 144, *N* = 72.

**Table S5.** The Post-hoc Model of Response Precision.

| Fixed Effects | Estimate | SE | *t* | 95% CI | *p* |  | Random Effects | Variance | SD |
| --- | --- | --- | --- | --- | --- | --- | --- | --- | --- |
| Intercept (Orientation) | 0.74 | 0.02 | 30.66 | [0.69, 0.78] | 0.000 |  | Subject (Intercept) | < 0.01 | 0.06 |
| Task (Location) | -0.34 | 0.03 | -10.53 | [-0.40, -0.28] | 0.000 |  |  |  |  |
| PC*_BULK_* | 0.02 | 0.01 | 2.25 | [0.00, 0.03] | 0.028 |  |  |  |  |
| PC*_DIR,_*_1_ | 0.02 | 0.01 | 1.50 | [0.00, 0.03] | 0.138 |  |  |  |  |
| PC*_DIR,_*_3_ | 0.02 | 0.02 | 1.30 | [-0.01, 0.05] | 0.200 |  |  |  |  |
| PC*_DIR,_*_1_ × PC*_DIR,_*_3_ | 0.04 | 0.01 | 3.35 | [0.02, 0.06] | 0.001 |  |  |  |  |
| PC*_DIR,_*_1_ × PC*_DIR,_*_3_ × Task (Location) | -0.03 | 0.02 | -1.65 | [-0.06, 0.00] | 0.103 |  |  |  |  |

**Table S6.** Model Comparison: The Best-fitting Model *vs.* Post-hoc Model of Response Precision.

| Model | Nested Model | Fixed Effects Added | Model Fit (ML) | | | | |  | LRT Against Nested | | |
| --- | --- | --- | --- | --- | --- | --- | --- | --- | --- | --- | --- |
|  |  |  | df | AIC | BIC | LL | Deviance |  | χ^2^ | df | *p* |
| *Best-fitting* | - | - | 8 | -38.51 | -14.76 | 27.26 | -54.51 |  |  |  |  |
| *Post-hoc* | *Best-fitting* | PC*_DIR,_*_1_ × PC*_DIR,_*_3_ × Task | 9 | -39.27 | -12.54 | 28.63 | -57.27 |  | 2.75 | 1 | 0.097 |

*Notes*: Total observations = 144, *N* = 72.

**Table S7.** The Post-hoc Model of Swap Errors.

| Fixed Effects | Estimate | SE | *t* | 95% CI | *p* |  | Random Effects | Variance | SD |
| --- | --- | --- | --- | --- | --- | --- | --- | --- | --- |
| Intercept (Orientation) | 0.03 | 0.01 | 3.31 | [0.01, 0.05] | 0.001 |  | Subject (Intercept) | < 0.01 | 0.03 |
| Task (Location) | 0.31 | 0.01 | 27.86 | [0.29, 0.33] | 0.000 |  |  |  |  |
| PC*_DIR,_*_1_ | 0.01 | 0.00 | 1.42 | [0.00, 0.01] | 0.162 |  |  |  |  |
| PC*_DIR,_*_3_ | 0.01 | 0.01 | 1.77 | [0.00, 0.02] | 0.082 |  |  |  |  |
| PC*_DIR,_*_1_ × PC*_DIR,_*_3_ | 0.01 | 0.00 | 1.96 | [0.00, 0.02] | 0.053 |  |  |  |  |
| PC*_DIR,_*_1_ × PC*_DIR,_*_3_ × Task (Location) | 0.00 | 0.01 | -0.10 | [-0.01, 0.01] | 0.919 |  |  |  |  |

**Table S8.** Model Comparison: The Best-fitting Model *vs.* Post-hoc Model of Swap Errors.

| Model | Nested Model | Fixed Effects Added | Model Fit (ML) | | | | |  | LRT Against Nested | | |
| --- | --- | --- | --- | --- | --- | --- | --- | --- | --- | --- | --- |
|  |  |  | df | AIC | BIC | LL | Deviance |  | χ^2^ | df | *p* |
| *Best-fitting* | - | - | 7 | -337.01 | -316.23 | 175.51 | -351.01 |  |  |  |  |
| *Post-hoc* | *Best-fitting* | PC*_DIR,_*_1_ × PC*_DIR,_*_3_ × Task | 8 | -335.02 | -311.27 | 175.51 | -351.02 |  | 0.011 | 1 | 0.917 |

*Notes*: Total observations = 144, *N* = 72.
